# Genetic diversity in the IIS6 domain of Voltage Gated Sodium Channel (*VGSC*) gene among *Aedes aegypti* populations from different geographical regions in India

**DOI:** 10.1101/2024.12.04.626773

**Authors:** MK Sumitha, G Navaneetha Pandiyan, M Kalimuthu, R Paramasivan, Bhavna Gupta

## Abstract

This study provides critical insights into the genetic diversity of the IIS6 domain of the *VGSC* gene in *Aedes aegypti* populations across various regions in India, focusing on two mutations: S989P and V1016G. Samples were collected from seven different cities across the country, including Dibrugarh, Kolkata, Berhampur, Bhopal, Bengaluru, Ghaziabad, and Aurangabad. The IIS6 domain was amplified and sequenced, revealing that the V1016G mutation was found at a higher frequency compared to the S989P mutation. The S989P mutation was most prevalent in Berhampur, followed by Bengaluru, while V1016G mutation showed high frequencies in Dibrugarh, followed by Berhampur. Additionally, the study identified intron polymorphisms within the *VGSC* gene, with the type A intron being relatively rare. However, the type A intron was observed in samples harbouring both mutant and wild alleles for both mutations. The regional variation in the frequencies of these mutations indicates complex evolutionary dynamics potentially influenced by local environmental factors and insecticide application practices. Interestingly, the high frequency of these alleles also correlated with the genetic structure of the mosquito populations, suggesting that gene flow might be playing a role in spreading these mutations. Regular monitoring of these mutations could serve as important indicators in assessing the status of resistance to pyrethroids and guide nationwide mosquito control efforts. This research underscores the necessity for localized vector control strategies and continuous genetic surveillance to manage insecticide resistance effectively.

## Introduction

*Aedes aegypti* is a primary vector for several significant arboviral diseases, such as dengue, chikungunya, Zika, etc. These diseases pose substantial public health challenges globally, particularly in tropical and subtropical regions. In India, dengue and chikungunya have caused numerous outbreaks (1–6), and Zika infections have recently emerged in several regions of the country (7,8) posing a significant burden on the healthcare system.

Effective control of mosquito populations is crucial in managing the spread of these vector-borne diseases. Chemical insecticides, including several pyrethroids such as deltamethrin, permethrin, cypermethrin, cyfluthrin, lambda-cyhalothrin, are commonly used for this purpose. However, the efficacy of these insecticides has been compromised due to the development of resistance in mosquito populations (9–12). A key mechanism of this resistance is associated with mutations in the voltage-gated sodium channel (*VGSC*) gene, also known as the knockdown resistance (*kdr*) mutations (13–15). Mutations in this gene can alter the target site of pyrethroids, reducing the insecticide’s effectiveness and leading to control failure (16,17). A number of *kdr* mutations such as V410L, G923V, L982W, S989P, A1007G, I1011M, V1016G, F1534C, D1763Y, associated with insecticide resistance have been identified worldwide (18–21). Of which S989P, V1016G, F1534C and I1011M are widely prevalent and have shown strong correlation with decreasing susceptibility to insecticides both in the field as well as in biophysiological experiments (22–28).

Voltage gated sodium channel (*VGSC*) gene is approximately 480 kb long encompassing 4 domains. The second domain (IIS6) of the *VGSC* gene encompasses important mutations like S989P, and V1016G/I that have been linked to pyrethroid resistance worldwide (18,21,29). Srisawat et al, identified S989P and V1016G mutations as probable candidates responsible for the development of deltamethrin resistance in *Ae. aegypti* Khu Bua strain (21). Furthermore, a study conducted in China reported the positive correlation of S989P and V1016G mutations to the resistance against cypermethrin and cyhalothrin (30). The coexistence of S989P and V1016G was first documented by Srisawat et al. in the Khu Bua strain of deltamethrin-resistant *Ae. aegypti*. S989P is hypothesized to synergize with V1016G when they coexist (21). Specifically, S989P is reported to reduce deltamethrin sensitivity ten-fold in combination with V1016G, while V1016G alone results in a two-fold reduction, and S989P by itself has no effect (Hirata et al., 2014).

India’s diverse environmental conditions and extensive use of insecticides create a complex landscape for the evolution of insecticide resistance in *Ae. aegypti* populations. Chemical use for mosquito control is very common, especially in urban areas. Municipalities regularly conduct larval control, and people use several chemical based personal protection measures (mosquito repellent liquidators, coils, sprays etc). Moreover, the country is also engaged in control and elimination programs for several vector-borne diseases like malaria, filariasis, and leishmaniasis, where vector control is the primary goal (National Strategic Plan: Malaria Elimination https://ncvbdc.mohfw.gov.in/Doc/National-Strategic-Plan-Malaria-2023-27.pdf). For example, insecticide-treated bed nets are heavily used in malaria endemic areas, such as in Odisha (31–33). The vector’s tolerance to different classes of insecticides continues to increase over the years as indicated by recent reviews (34,35). The status of insecticide resistance among *Ae. aegypti* populations in India is a pressing concern, with diverse degrees of resistance observed across different regions (34,36–38).

Therefore, regular screening of mosquito populations is crucial to monitor resistance status. Genotyping of insecticide resistance mutations could serve as a valuable surveillance tool to reveal patterns of resistance that may be unique to specific regions or widespread across the country. Changes in mutation frequency over the years can be correlated with insecticide pressure, providing a basis for tailoring local vector control programs and informing national public health policies. However, such studies on Indian *Ae. aegypti* populations are scarce and are restricted to single population (28,39–42). To fill this gap, this study was carried out to genotype the IIS6 domain of the *VGSC* gene in several *Ae. aegypti* populations from India. Overall, this study has revealed the current situation of insecticide resistance mutations and has also generated a reference base for future comparisons.

## Material and methods

### Study area and sample collection

*Ae. aegypti* samples were collected from various locations across India, including Dibrugarh (Assam), Kolkata (West Bengal), Berhampur (Odisha), Bhopal (Madhya Pradesh), Bengaluru (Karnataka), Ghaziabad (Uttar Pradesh) and Aurangabad (Maharashtra) during 2019 and 2020 (Figure 1; Table 1). The samples were collected as both larval and adult collections. Larval samples were reared in the laboratory until emergence, ensuring that only a single mosquito from each larval container was used for this analysis. The collection methodology followed the procedures outlined in our previous studies (43–45). *Ae. aegypti* was identified based on distinctive morphological characteristics, including black and white markings on its legs and a lyre-shaped marking on the upper side of its thorax, as described in the morphological key written by Christophers (46). The number of samples and their respective locations are detailed in Table 1 and illustrated in Figure 1.

**Figure 1:**
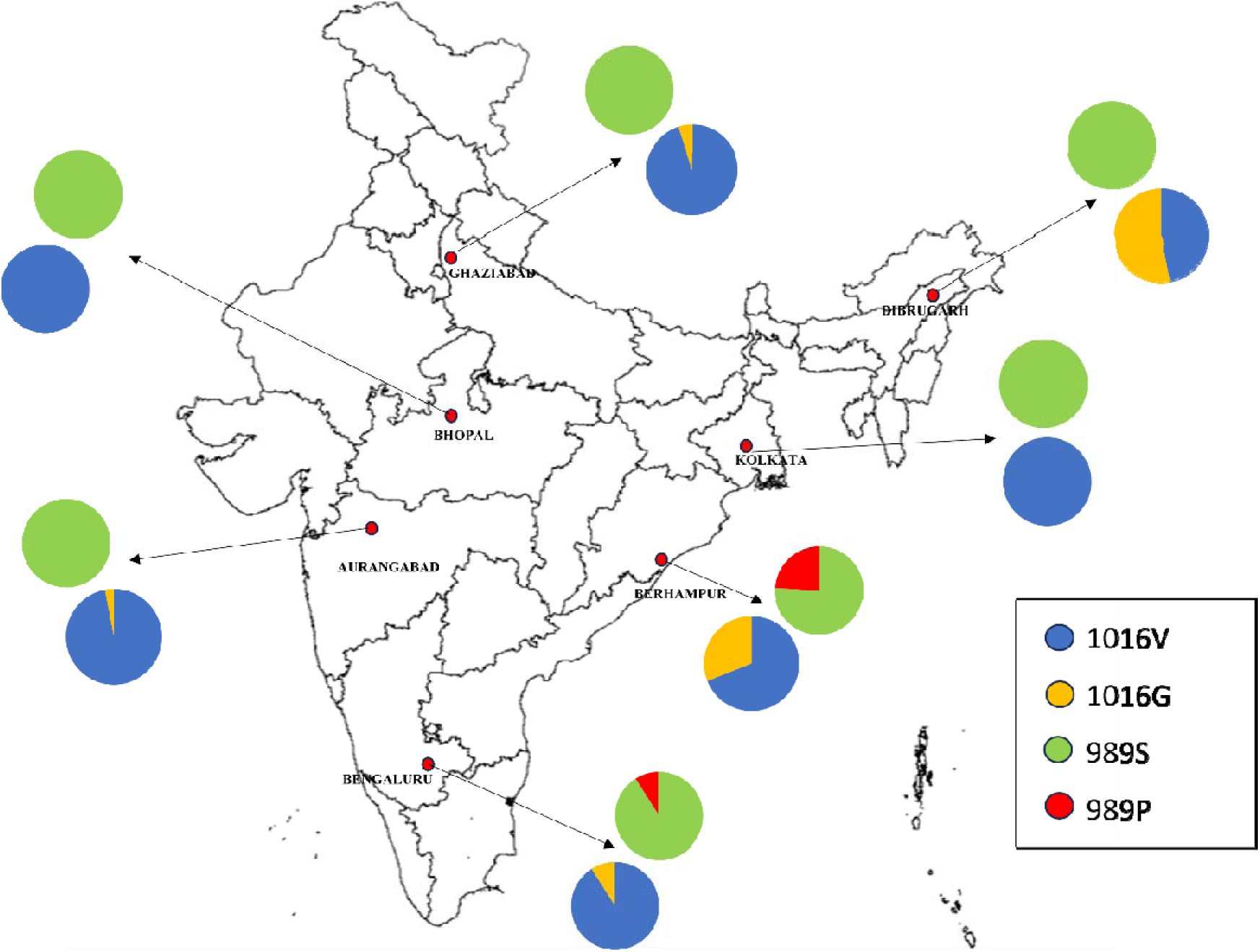
Sampling sites from India displaying the distribution of *kdr* mutations (S989P and V1016G) in *Ae. aegypti*. Pie charts represent the allelic frequencies.

**Table 1.** Location wise genotyping results of S989P and V1016G mutations identified through sequencing.

|  |  | Total samples<br>(140) | Dibrugarh<br>(17) | Aurangabad<br>(18) | Bengaluru<br>(22) | Bhopal<br>(22) | Ghaziabad<br>(11) | Kolkata<br>(19) | Berhampur<br>(31) |
| --- | --- | --- | --- | --- | --- | --- | --- | --- | --- |
| Genotypic<br>frequency | SS | 124 (88.6%) | 17 (100%) | 18 (100%) | 18 (81.81%) | 22 (100%) | 11 (100%) | 19 (100%) | 19 (61.30%) |
|  | SP | 13 (9.3%) | 0 | 0 | 4 (18.19%) | 0 | 0 | 0 | 9 (29.03%) |
|  | PP | 3 (2.1%) | 0 | 0 | 0 | 0 | 0 | 0 | 3 (9.67%) |
| Allelic<br>frequency | S | 0.93 | 1 | 1 | 0.91 | 1 | 1 | 1 | 0.76 |
|  | P | 0.07 | 0 | 0 | 0.09 | 0 | 0 | 0 | 0.24 |
| Genotypic<br>frequency | VV | 105 (75%) | 4 (23.53%) | 17 (94.44%) | 18 (81.82%) | 22 (100%) | 10 (90.91%) | 19 (100%) | 15 (48.39%) |
|  | VG | 27 (19.3%) | 8 (47.06%) | 1 (5.56%) | 4 (18.18%) | 0 | 1 (9.09%) | 0 | 13 (41.94%) |
|  | GG | 8 (5.7%) | 5 (29.41%) | 0 | 0 | 0 | 0 | 0 | 3 (9.67%) |
| Allelic<br>frequency | V | 0.85 | 0.47 | 0.97 | 0.91 | 1 | 0.95 | 1 | 0.69 |
|  | G | 0.15 | 0.53 | 0.03 | 0.09 | 0 | 0.05 | 0 | 0.31 |
| Intron type | A | 16 (11.43%) | 5 | 1 | 3 | 0 | 0 | 0 | 7 |
|  | B | 80 (57.14%) | 4 | 13 | 6 | 21 | 10 | 18 | 8 |
|  | Mixed | 44 (31.43%) | 8 | 4 | 13 | 1 | 1 | 1 | 16 |

### Amplification and sequencing of IIS6 domain of *VGSC* gene

The genomic DNA was isolated from individual mosquitoes using the DNeasy Blood and Tissue kit (QIAGEN) following the manufacturer’s instructions. Primers IIP_F (5′ -GGT GGA ACT TCA CCG ACT TC-3′) and IIS_R (5′-GGA CGC AAT CTG GCT TGT TA-3′) were used to amplify the partial sequence of domain II, covering regions that encompass codons encoding amino acid residues 989 and 1016 in the IIP-IIS6 region (47). DNA products of 581 bp were generated by PCR.

The reactions were carried out in a final volume of 20 µl, comprising 0.2 µl of Taq polymerase, 4 µl of 5X buffer, 2 µl of dNTP, 0.4 µl of 10 mM IIP_F and 0.4 µl of 10 mM of IIS_R, 100-150 ng of DNA template, and water to make up to 20 µl. The amplification consisted of an initial heat-activation step of 95°C for 3 minutes, followed by 35 cycles of: 95°C for 30 seconds, 60°C for 30 seconds, and 72°C for 30 seconds. Final extension was done at 72°C for 5 minutes. The PCR products were loaded into a 2% agarose gel and run for 45 minutes at 70V. The bands were visualized under UV light in a transilluminator. The target bands were delicately cut out, and the DNA fragments were extracted and purified using the Wizard® SV Gel and PCR Clean-Up System and were sequenced from both directions using forward (IIP_F) and reverse (IIS_R) primers.

### Sequence Analysis

The sequences of the fragments were analyzed using MEGA 11 (Molecular Evolutionary Genetic Analysis software) (48), aligned by Clustal W (49), and the single nucleotide polymorphisms and intron types were meticulously identified and documented. The sequences generated in this study have been submitted in the GenBank. Haplotype network using median joining algorithm was generated using Network 10.2.0.0 (50).

## Results

The IIS6 domain of the *VGSC* gene was sequenced in a total of 175 *Ae. aegypti* mosquitoes collected from various regions in India. However, upon sequence quality assessment, only 140 samples yielded high-quality sequences suitable for further analysis. The sequences of the IIS6 domain in 140 individual *Ae. aegypti* specimens revealed the presence of only two non-synonymous mutations: S989P and V1016G. No other previously identified *kdr* mutations, such as A1007G, I1011M, and L1014S, were observed. The mutation S989P was observed in 16 out of 140 samples (11%) and V1016G was observed in 35 samples (25%). The frequencies of each mutation varied across different regions (Table 1).

The S989P mutation was most prevalent in Berhampur, with a frequency of 0.24. Of these, three samples exhibited homozygous mutations, and nine samples had heterozygous mutations. This mutation was also detected in Bengaluru, where four samples had heterozygous mutations (Table 1). However, the mutation was absent in the other five regions sampled.

The V1016G mutation was more prevalent compared to S989P. The highest frequencies were observed in Dibrugarh (0.53), followed by Berhampur (0.31). It was not observed in Bhopal and Kolkata. In Dibrugarh, five samples had homozygous mutations and eight were heterozygous. In Berhampur, 13 samples had heterozygous mutations and three had homozygous mutations. The mutation was also found in heterozygous conditions in Aurangabad, Bengaluru and Ghaziabad in lower frequencies (Table 1).

In our analysis of 140 *Ae. aegypti* samples, the majority, comprising 105 samples (75%), exhibited the wild-type genotype for both the S989P and V1016G mutations. Notably, three samples from Berhampur were identified as double mutants, carrying homozygous mutations for both S989P and V1016G. Additionally, 13 samples, primarily from Berhampur (nine samples) with a smaller representation from Bengaluru (four samples), displayed a double heterozygous genotype for both mutations. Furthermore, 14 samples (eight from Dibrugarh, four from Berhampur and one each from Ghaziabad and Aurangabad) showed heterozygosity solely for the V1016G mutation, while five samples from Dibrugarh were homozygous for this mutation alone.

These observations led to the identification of five haplotypes based on the presence or absence of the S989P and V1016G mutations: SV (wild type), PG (double homozygote), SP/VG (double heterozygotes), SG (homozygote for wild S989 and mutant 1016G), and S/VG (heterozygote for V1016G with wild allele for S989) (Figure 2, Supplementary Figure 1). Of particular note was the consistent association of the 989P mutation with the 1016G mutation across all three samples.

**Figure 2:**
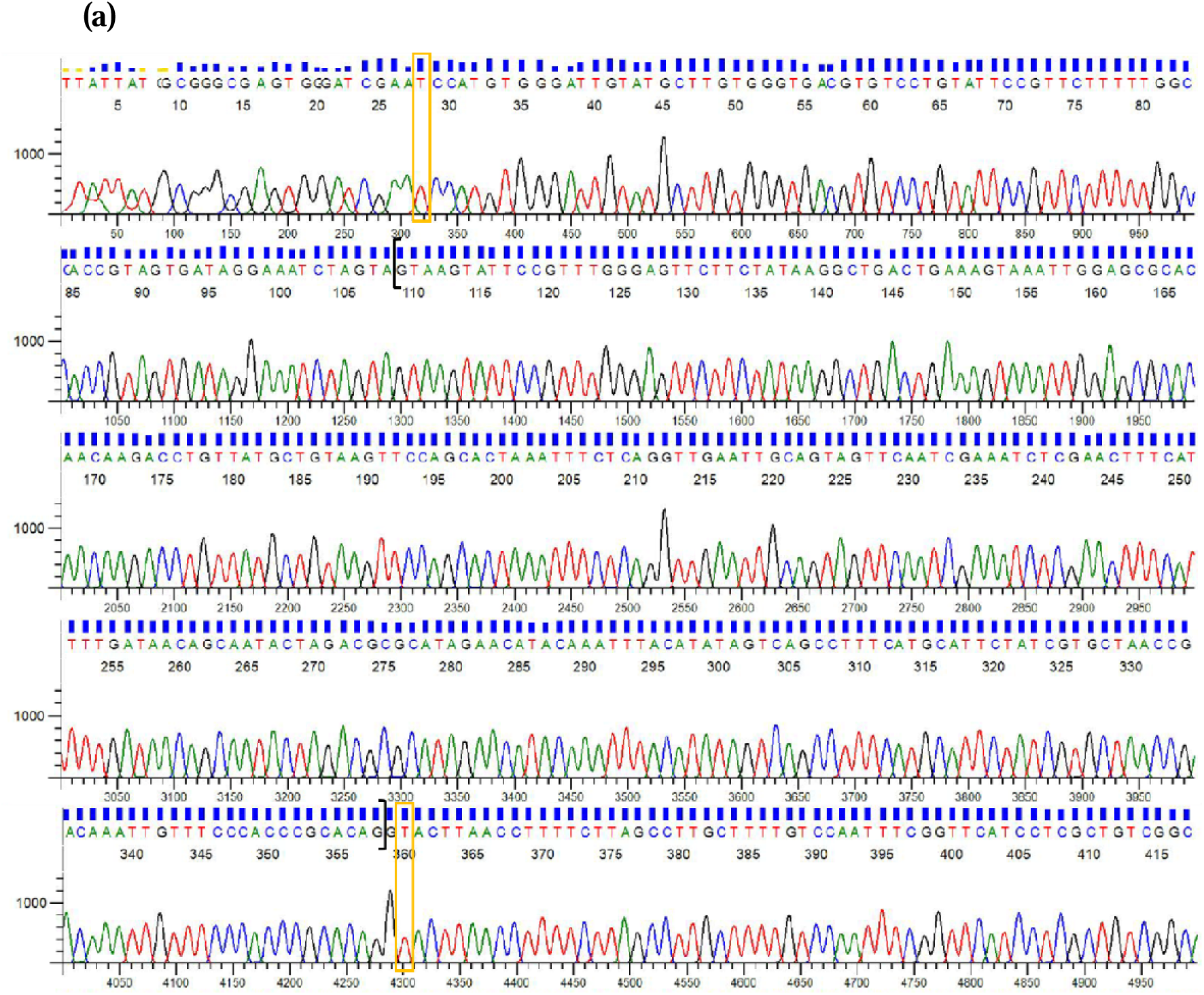
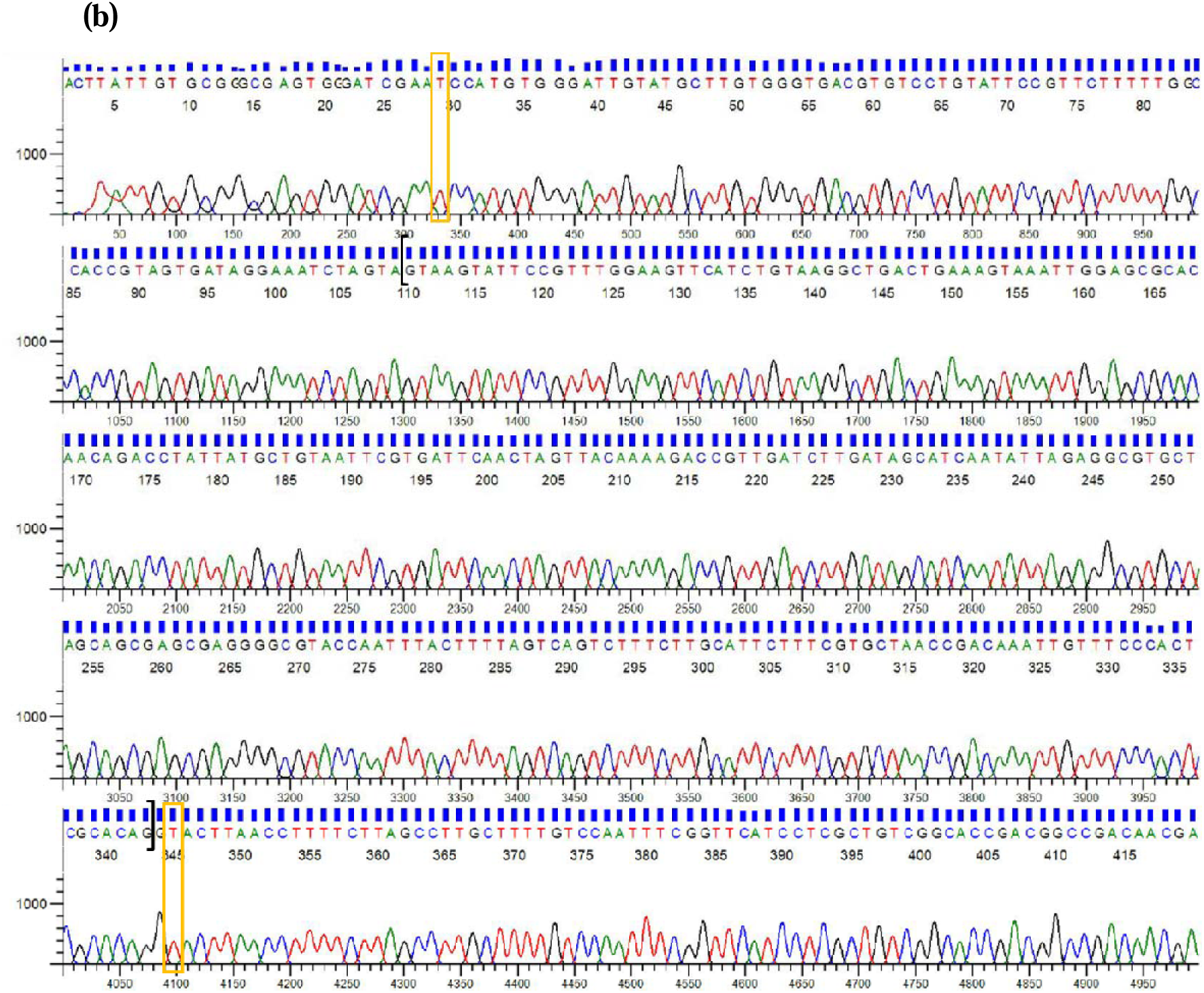
Chromatograms showing sequences of IIS6 segments of *Ae. aegypti* (a) wild type with type A intron and (b) wild type with type B intron. Yellow boxes represent the position of the mutations and the squares brackets enclose the introns.

The intron situated between exon 20 and 21 within the domain IIS6 exhibited notable polymorphisms. While detailed sequence analysis of the intron region was not conducted, variations in length were observed, indicating polymorphism. Specifically, two distinct intron types, designated as intron A (250bp) and intron B (234bp), were identified across the *Ae. aegypti* populations under study. Among the samples analyzed, 16 were characterized by the presence of intron A, while 80 exhibited intron B. Additionally, 44 samples displayed a mixed type of intron. The chromatograms showing wild type alleles with each intron type are shown in Figure 2.

Consistent with the previous findings (19,51), our analysis revealed that double homozygotes and homozygotes for 1016G mutation were consistently associated with the type A intron. Similarly, both double heterozygotes and single homozygotes for the V1016G mutation were observed in association with either the A type or a mixed type intron (Figure 3). V1016G heterozygotes however, had either B type or mixed introns. Moreover, wild-type samples also displayed variability in intron types, with six samples featuring the A type intron, 78 harbouring the B type intron, and 21 exhibiting a mixed type intron. To elucidate the relationships and associations among intron types, haplotypes, and observed mutations, a haplotype network was constructed (Figure 3).

**Figure 3:**
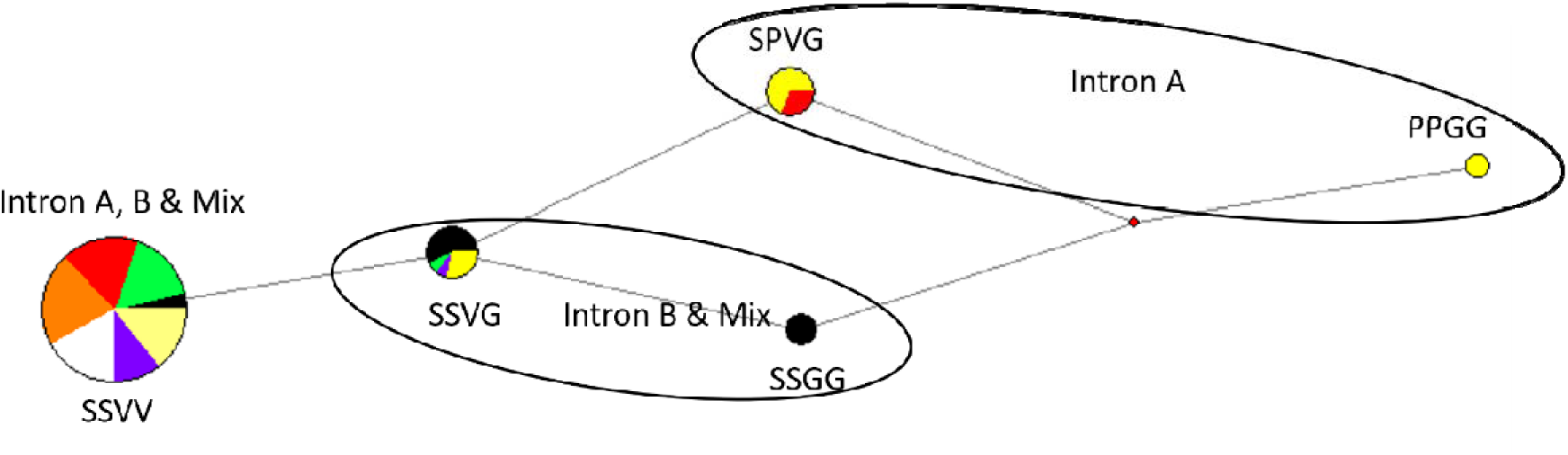
Haplotype network based on the S989P and V1016G mutations, showing the association with intron types A and B. The network highlights the distribution and frequency of each haplotype within the sampled populations.

## Discussion

### Trends in *kdr* mutations and regional variations

This study revealed the presence of two non-synonymous mutations, S989P and V1016G, in the IIS6 domain of the *VGSC* gene of *Ae. aegypti* across different regions of India. These mutations, resulting from nucleotide substitutions TCCLJ→LJCCC and GTALJ→LJGGA, respectively, were found at varying frequencies among different regions. The absence of other known *kdr* mutations, such as A1007G, I1011M, and L1014S, suggests regional specificity in the mutation profiles of *Ae. aegypti* populations in India.

Previous studies conducted in various regions of India have reported the presence of S989P and V1016G. For instance, research conducted in Bengaluru observed high frequencies of these mutations (0.18 S989P and 0.18 V1016G) and revealed a low-level association between *kdr* genotypes and resistance to DDT, deltamethrin, and permethrin (28). A study in Delhi in 2015 could not identify S989P and V1016G, but reported F1534C and T1520I in *VGSC* gene among *Ae. aegypti* populations resistant to DDT, permethrin, and deltamethrin (40). Additionally, high frequencies of S989P and V1016G were observed in Punjab (39) and West Bengal (42) among populations exhibiting insecticide resistance. In our study, S989P was detected with a frequency of 0.07 while V1016G was observed with a frequency of 0.15. However, these frequencies varied across different geographical regions, underscoring the considerable variability in mutation presence and resistance patterns observed across distinct regions of India.

The majority of the samples (75%) were found to be wild type, with no mutations detected. The samples from Bhopal and Kolkata did not possess any of the observed mutations. Furthermore, samples from Aurangabad and Ghaziabad exhibited low frequencies of mutant alleles, with only 0.03 and 0.05 possessing the V1016G mutation, respectively. This variation in mutation frequencies across different regions may be attributed to varying levels of insecticide pressure on mosquito populations or the absence of strong associations between these mutations and resistance, which has not been confirmed by bioassays in this study. However, the high frequencies of these mutations observed in Berhampur, Dibrugarh, and Bengaluru raise concerns and suggest their potential evolutionary or biological significance in the context of insecticide resistance in *Ae. aegypti* populations.

The regional differences in frequencies might be due to different insecticide usage patterns, environmental conditions, and mosquito population dynamics. For example, high level of diversity in both the mutations were observed in Berhampur in Odisha. Odisha, one of the most malaria-endemic regions in India, has implemented extensive mosquito control activities, including the use of long-lasting insecticide-impregnated nets under malaria elimination program. Moreover, several mosquito control programs have been carried out in Odisha. These measures may exert significant selection pressure on mosquito populations, potentially leading to the high frequency of *kdr* mutations.

It is also interesting to note that our recent population genetic analysis of *Ae. aegypti* across India revealed that populations from Dibrugarh, Berhampur, Kolkata, Bhopal, and Bengaluru clustered together (Sumitha et al., 2023a). While *kdr* mutations were not observed in Bhopal and Kolkata, the mutations were present in the other three regions. The shared genetic characteristics among these regions or different selection pressures due to variations in insecticide use patterns might explain the higher frequency of *kdr* mutations compared to other regions in India, where these mutations are less prevalent. However, the regional variations in the frequency of these mutations suggests the need for targeted vector control strategies tailored to specific regional conditions to effectively manage and mitigate the spread of insecticide resistance.

### Linkage disequilibrium and mutation synergy

Previous studies have reported complete linkage disequilibrium between S989P and V1016G mutations (28,51–54), indicating that these mutations tend to co-occur more frequently than expected by chance alone (Kawada et al., 2016). For example, research conducted in Bengaluru and Saudi Arabia demonstrated complete linkage disequilibrium between these two mutations, suggesting a strong association between them. However, our study observed only partial linkage disequilibrium, as the S989P allele was consistently found in association with the V1016G mutation, either in homozygous or heterozygous conditions. However, five samples from Dibrugarh having homozygous V1016G mutations were also found in isolation. This discrepancy in linkage patterns across different populations could be attributed to various factors such as differences in selection pressures exerted by insecticide usage, genetic drift, or gene flow between populations. It’s possible that in regions where insecticide usage is more intense or where specific insecticides are more frequently used, complete linkage disequilibrium may be observed due to strong selection pressures favouring the co-occurrence of these resistance-conferring mutations. On the other hand, populations experiencing lower selection pressures or undergoing genetic drift may exhibit partial linkage disequilibrium or even the presence of isolated mutations.

The presence of mutations is also influenced by their biological significance. It has been observed that S989P reduces deltamethrin sensitivity by ten-fold in combination with V1016G, whereas V1016G alone results in a two-fold reduction in deltamethrin sensitivity, while S989P has no effect on its own (55). Thus, understanding the dynamics of linkage disequilibrium between *kdr* mutations can provide valuable insights into the evolution and spread of insecticide resistance in *Ae. aegypti* populations. Further research incorporating broader geographic sampling and detailed phenotypic characterization through bioassays would be beneficial to elucidate the complex interplay between *kdr* mutations and their contribution to insecticide resistance.

### Intron polymorphisms

The two intron types; intron A (250 bp) and B (234 bp) that have been widely observed in *Ae. aegypti* populations (56) were recorded in our samples. This intron is located between the amino acid residues 1015 and 1016 (19). The type A intron is found strongly associated with the V1016G mutation (51) as well as the V1016I mutation (19). In our samples, type A intron was observed in all the samples having either V1016G or both the mutations. This intron type was also observed in samples with wild type alleles. Berhampur had the highest number of type A intron, followed by Dibrugarh, Bengaluru, Aurangabad and Ghaziabad. 16 samples including seven from Berhampur, five from Dibrugarh, three from Bengaluru and one from Aurangabad had only A type, and 21 samples had mixed introns (both A & B). The presence of different intron types might influence the splicing and expression of the *VGSC* gene, potentially affecting resistance mechanisms. Further studies could investigate the functional implications of these intron polymorphisms and their role in the evolution of insecticide resistance. The presence of mixed introns samples suggests a high level of genetic diversity within these populations. This genetic diversity could be a result of gene flow between populations or the presence of heterozygous individuals, which can harbour both intron types. Such diversity might provide a survival advantage to mosquito populations by maintaining a pool of genetic variants that can be selected under different environmental conditions and insecticide pressures.

### Limitations and future directions

The findings from this study provide valuable insights into the genetic diversity and distribution of *kdr* mutations in *Ae. aegypti* populations across India. However, the absence of bioassay data limits our ability to definitively link these mutations to insecticide resistance. The observed regional variations in mutation frequencies underscore the need for localized vector control strategies. For instance, high diversity in this domain was observed in Berhampur in Odisha, and Dibrugarh in Assam. Both regions are endemic to malaria and undergoing extensive mosquito control activities under malaria and filariasis elimination programs. In these areas, the replacement or modification of current insecticide regimens may be considered. Moreover, continuous monitoring and genetic analysis of mosquito populations in these regions are crucial for adapting more effective control measures and mitigating the spread of resistance. Future studies could complement genetic analysis with bioassays to provide a more comprehensive understanding of insecticide resistance dynamics. Integrating molecular data with field surveillance and resistance monitoring will be crucial in developing sustainable and adaptive vector management programs.

## Acknowledgements

Authors would like to thank Indian Council of Medical Research (ICMR) for the intramural facilities to carry out this study. MK Sumitha and G Navaneetha Pandiyan would like to thank Madurai Kamaraj University for supporting their research.

---

**Supplementary Figure 1:**
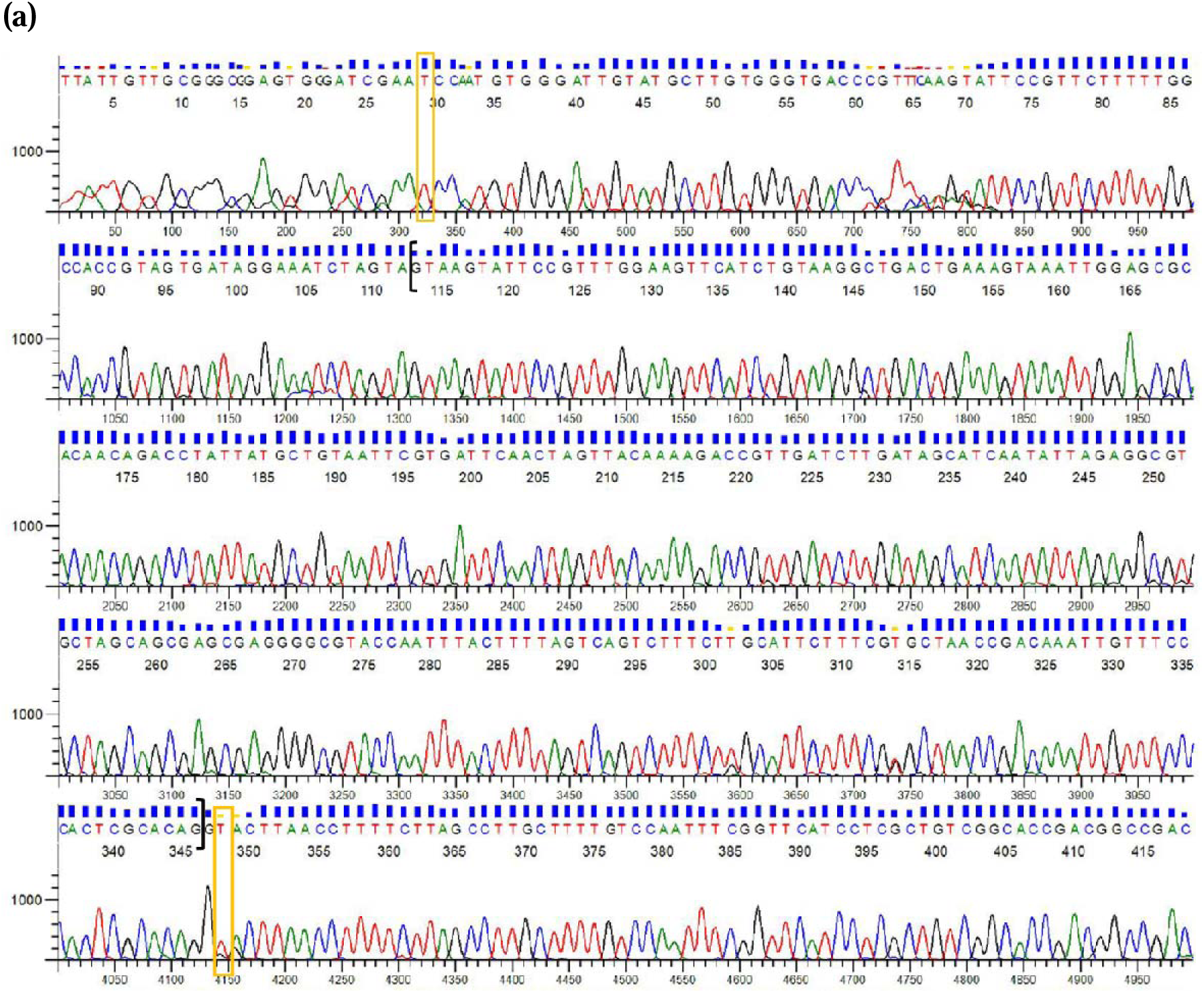
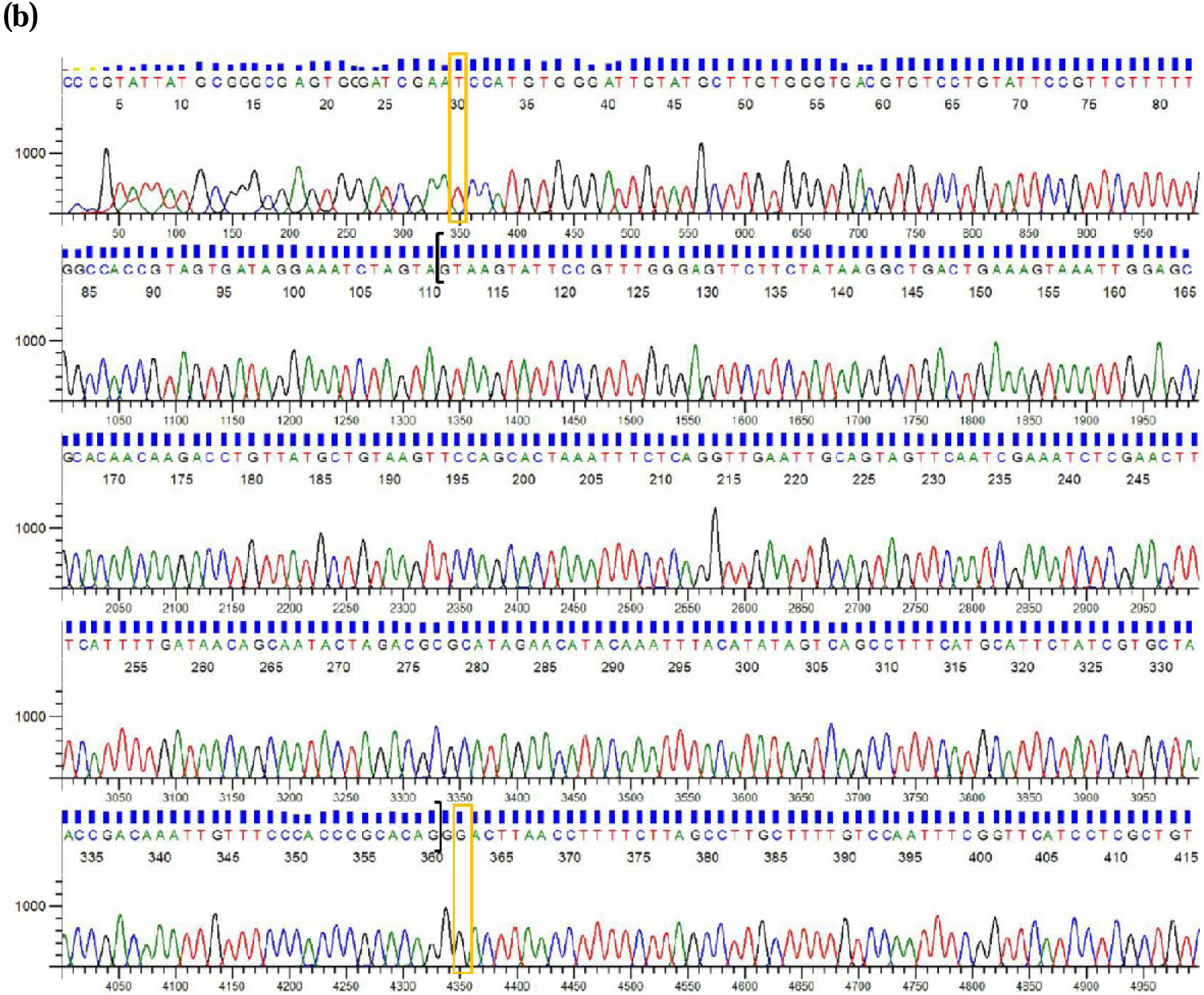
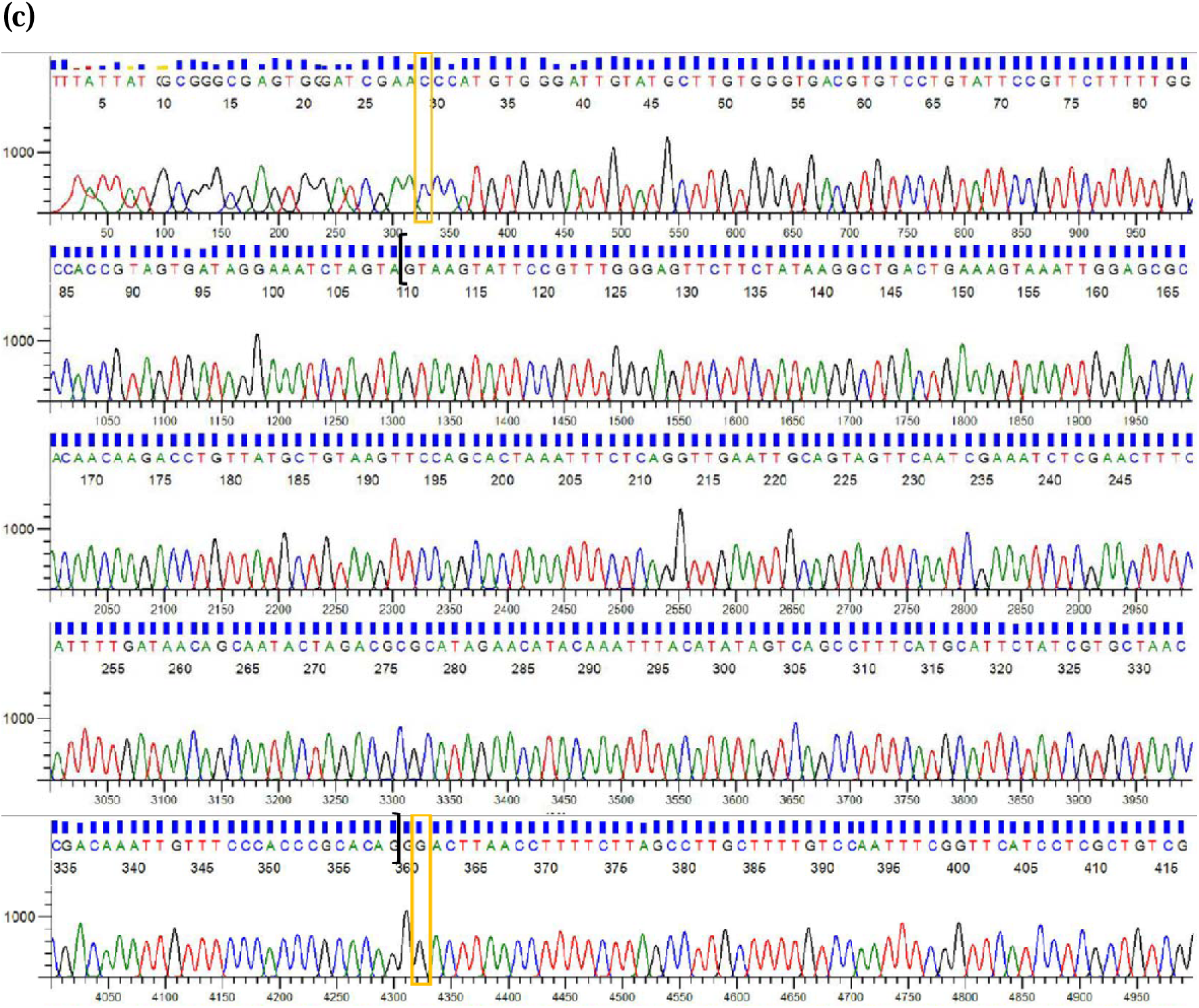
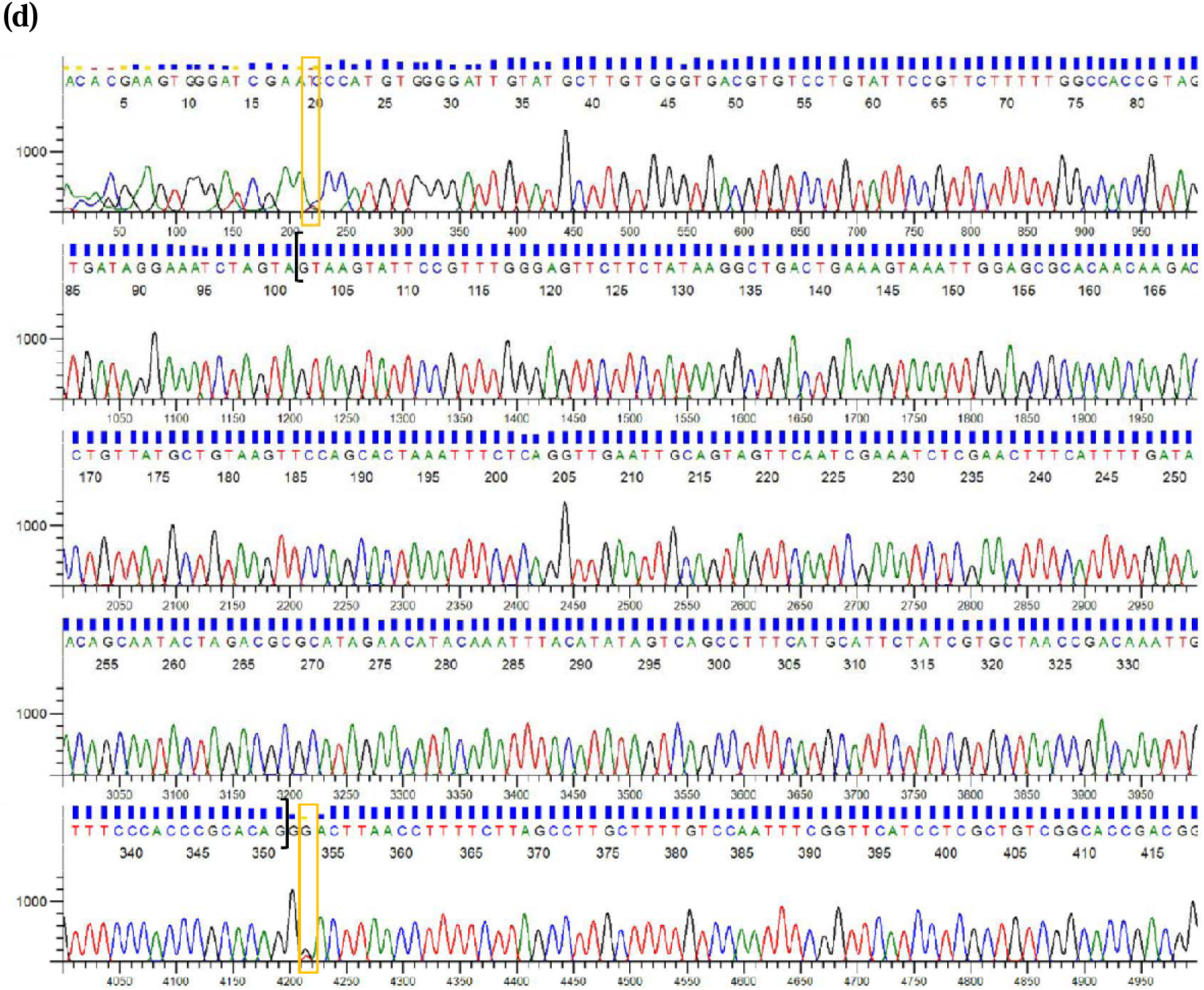
Chromatograms showing sequences of IIS6 segments of *Ae. aegypti* (a) heterozygous mutant for V1016G with type B intron, (b) homozygous mutant for 1016G with type A intron, (c) double homozygous mutant for 989P and 1016G with type A intron and (d) double heterozygous mutant with type A intron. Yellow boxes represent the position of the mutations and the squares brackets enclose the introns.

