## Supplementary figures and images for "Genetic diversity in the IIS6 domain of Voltage Gated Sodium Channel (*VGSC*) gene among *Aedes aegypti* populations from different geographical regions in India"

### Supplementary figure 1a

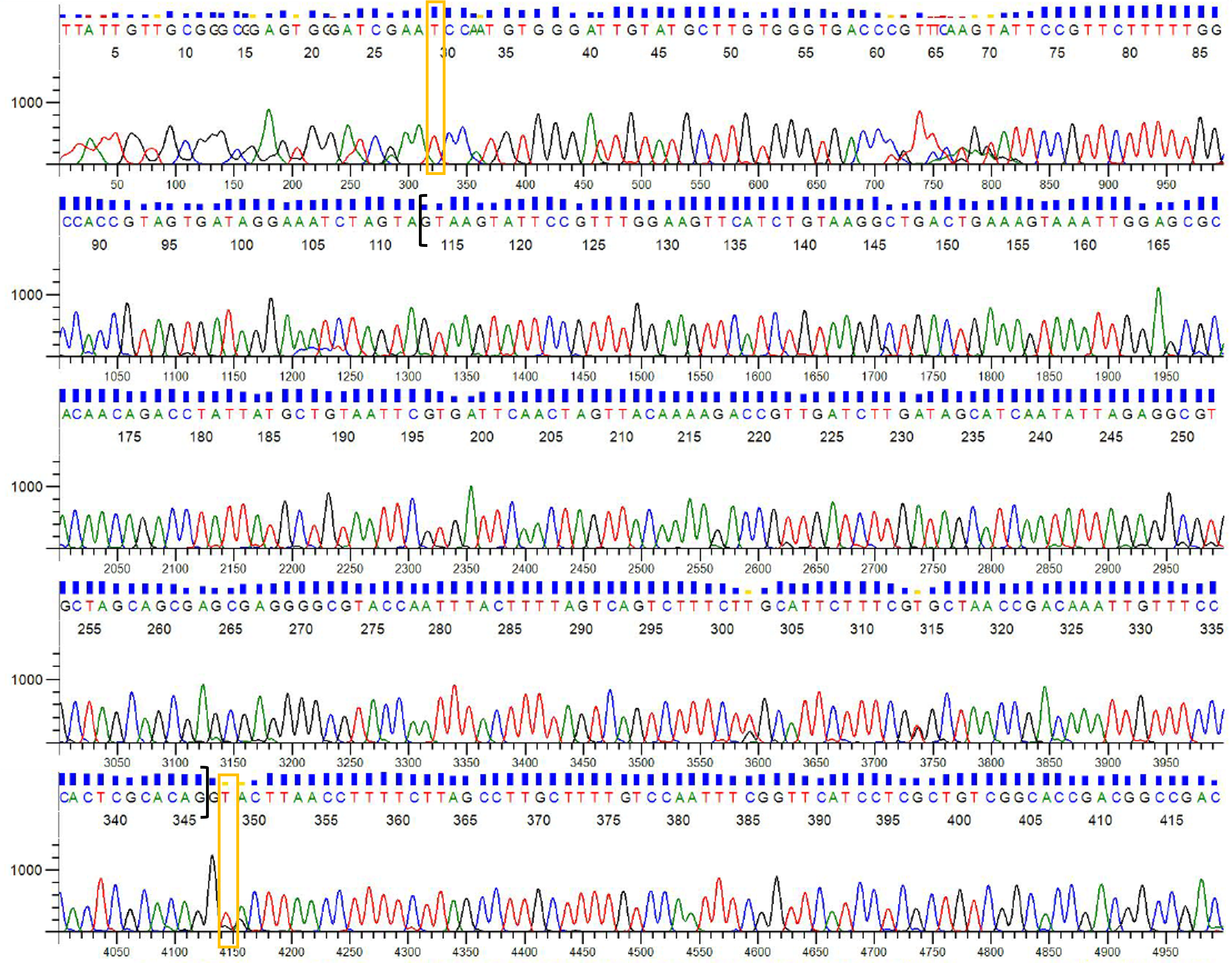

### Supplementary figure 1b

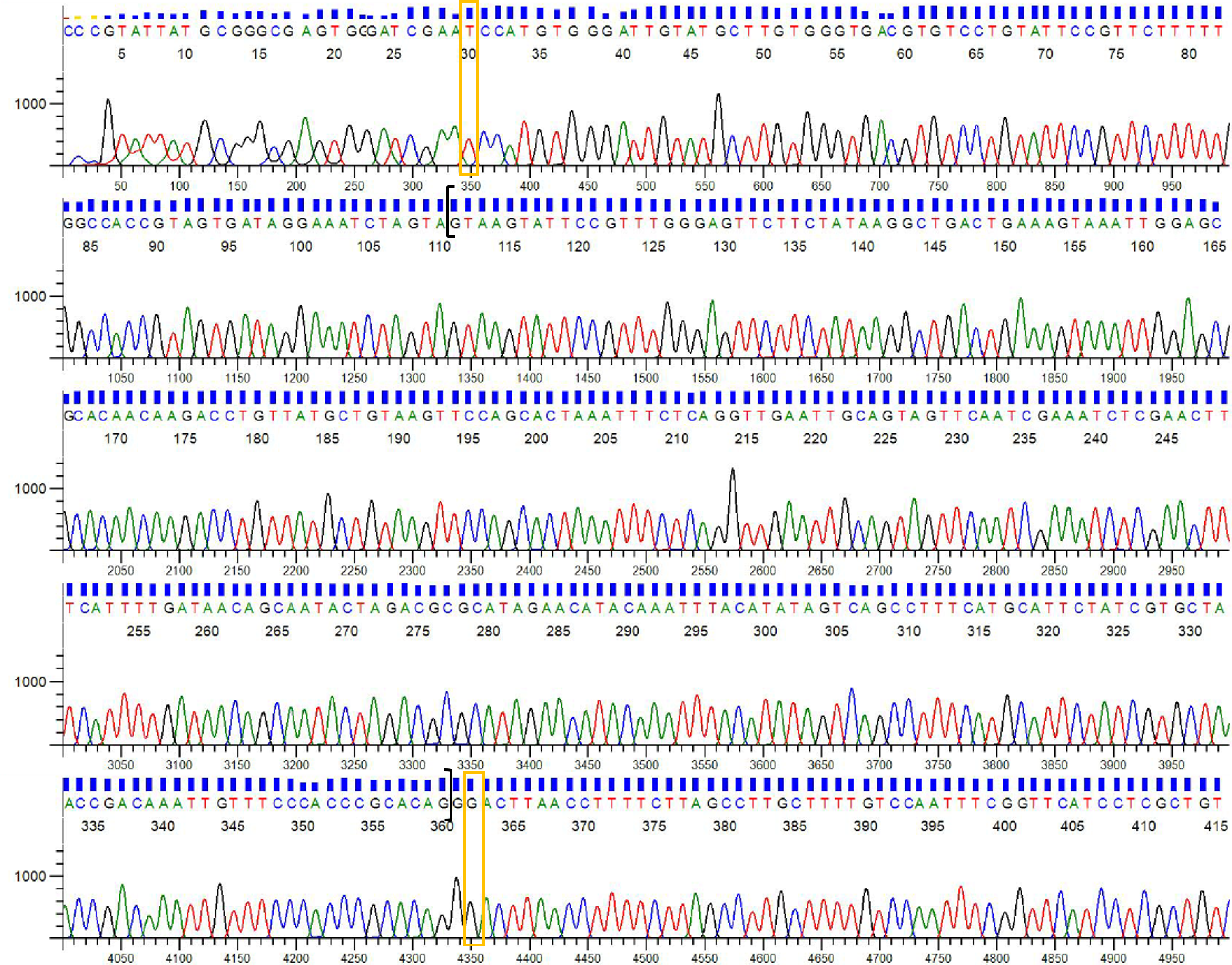

### Supplementary figure 1c

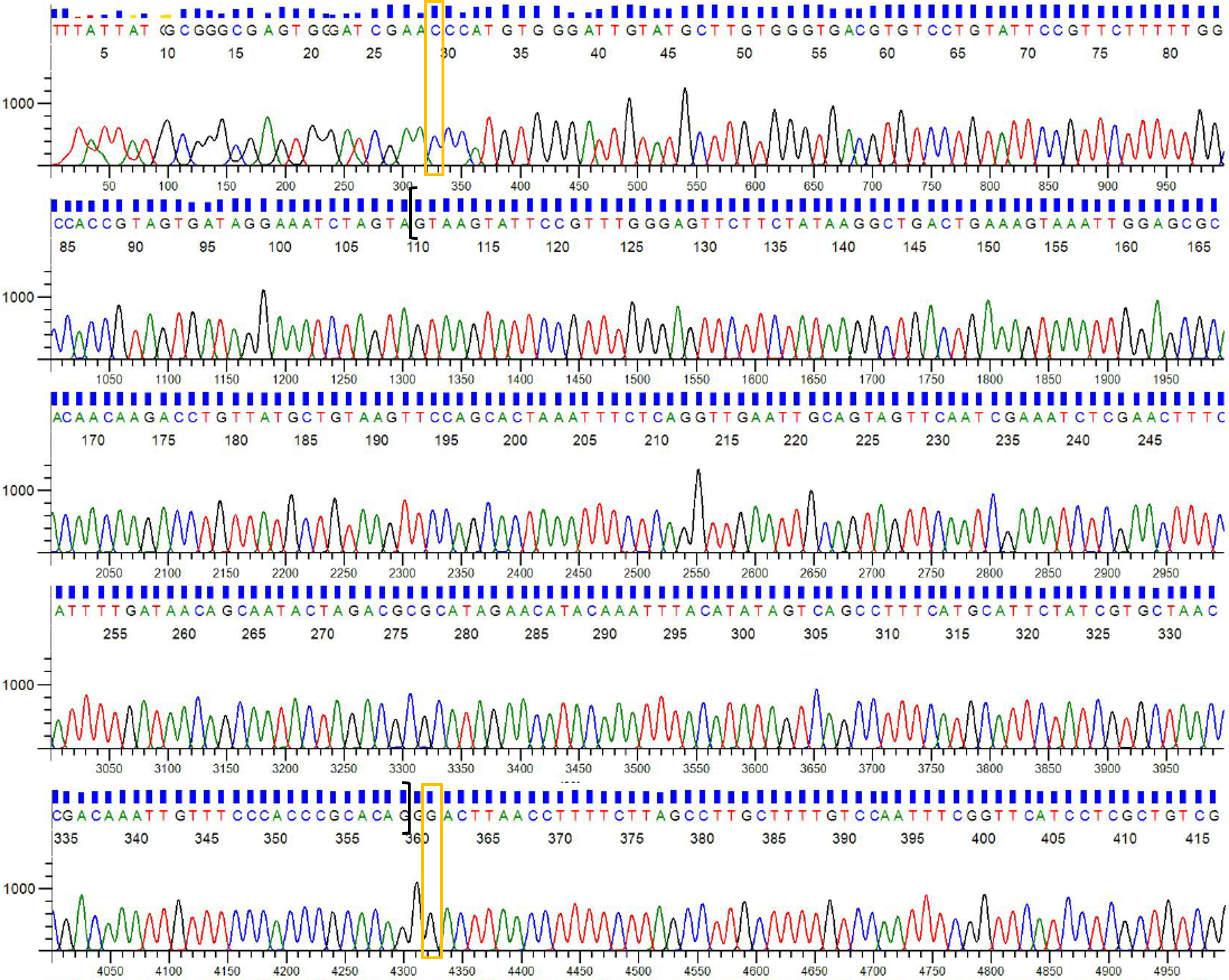

### Supplementary figure 1d

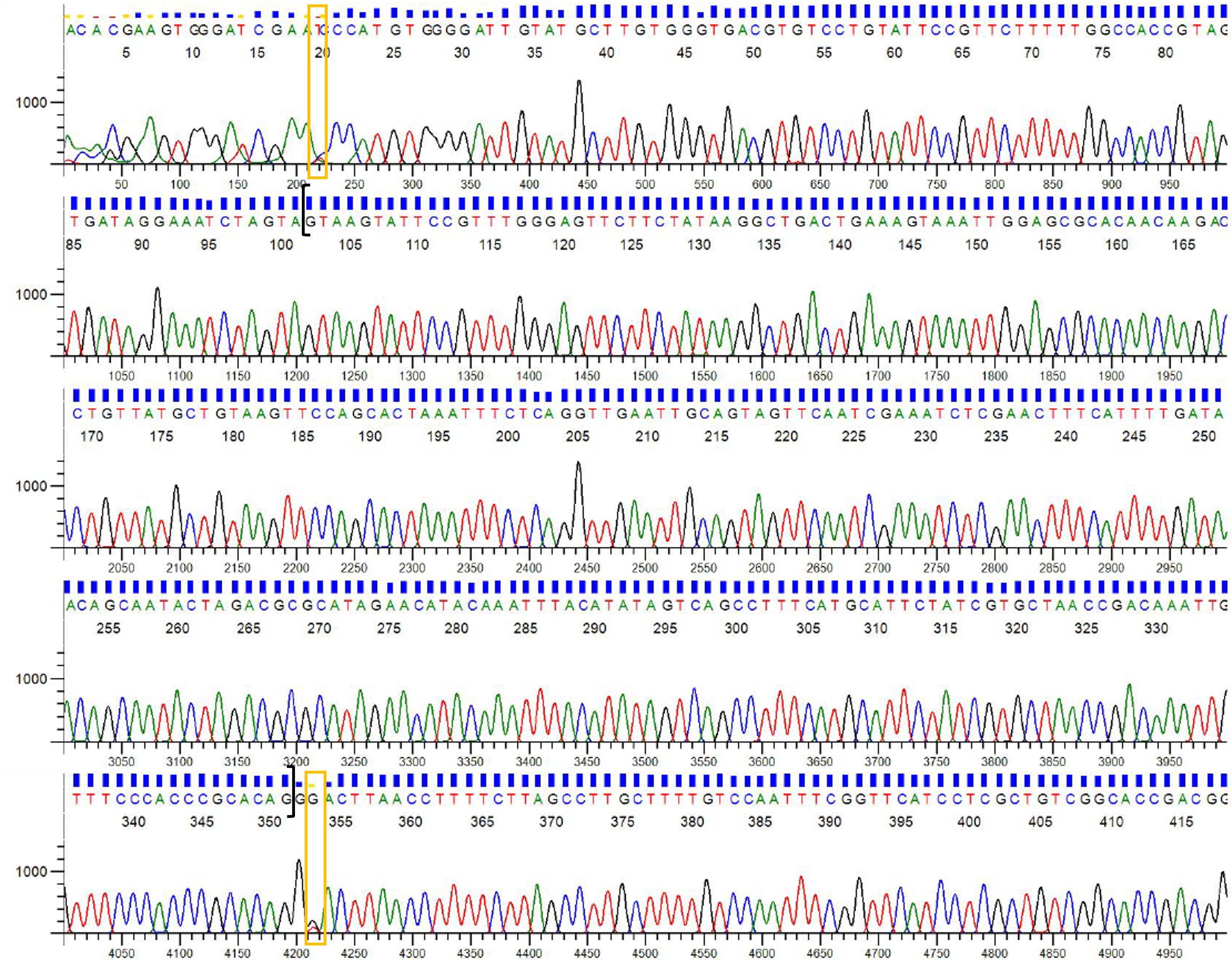
